# Structural insight into hormone recognition and transmembrane signaling by the atrial natriuretic peptide receptor

**DOI:** 10.1101/2020.07.09.196394

**Authors:** Haruo Ogawa, Masami Kodama, Kei Izumikawa

## Abstract

Atrial natriuretic peptide (ANP) is an endogenous and potent hypotensive hormone that elicits natriuretic, diuretic, vasorelaxant, and anti-proliferative effects, which play a central role in the regulation of blood pressure and volume. To investigate the hormone-binding and membrane signaling mechanisms mediated by the ANP receptor, we elucidated the crystal structures of the ANP receptor extracellular hormone-binding domain (ANPR) complexed with full-length ANP, truncated mutants of ANP, and dendroaspis natriuretic peptide (DNP) isolated from the venom of the green Mamba snake, *Dendroaspis angusticeps*. The bound peptides possessed pseudo two-fold symmetry to which the tight coupling of the peptide to the receptor and guanylyl cyclase activity was attributed. The crystal structures and our results from kinetic experiments provide insight into the ligand recognition and transmembrane signaling mechanism of the ANP receptor. Our findings provide useful information that can be applied to drug discovery for heart failure therapies.

## Introduction

The cardiac hormone, atrial natriuretic peptide (ANP)^1^,is produced in the cardiac atria and secreted into the circulatory system in response to volume expansion and increased atrial distension. ANP has potent natriuretic, diuretic, vasodilator, and renin- and aldosterone-suppressing activities, which plays a major role in the regulation of blood pressure and volume regulation^2, 3, 4^. Transgenic mice overexpressing ANP have been shown to have reduced blood pressure^5^. ANP gene knockout mice have been shown to develop salt-sensitivity^6^, while ANP receptor gene knockout mice developed salt-insensitive hypertension^7^, ventricular hypertrophy, and sudden death^8^. These findings indicate that the signaling pathways mediated by ANP and ANP receptors are indispensable for cardiovascular pathophysiology. ANP has also been reported to prevent cancer metastasis through the repression of inflammatory reactions^9^. ANP is a cyclic peptide of 28 amino acid residues^10^ and belongs to the natriuretic peptide family, which includes B-type (BNP)^11^, and C-type (CNP)^12^ natriuretic peptides. The activities of BNP and ANP are similar^13^, whereas CNP is thought to play roles in the central nervous system-mediated control of blood pressure and salt-fluid balance^13, 14, 15^, cartilage homeostasis, and endochondral bone formation^16^. Dendroaspis natriuretic peptide (DNP) has recently been isolated from the venom of the green Mamba, *Dendroaspis angusticeps*^17^. DNP is a potent natriuretic and diuretic peptide, similar to ANP and BNP, which elicits increases in urinary and plasma cGMP^18, 19^.

The activities of ANP and BNP are mediated by the ANP receptor, a single-span transmembrane receptor harboring intrinsic guanylate cyclase (GCase) activity^20^. The ANP receptor consists of a glycosylated extracellular hormone-binding domain and an intracellular domain that includes protein kinase-like and GCase catalytic domains, and acts as a homodimer. The closely related B-type natriuretic (GC-B) receptor mediates the activities of CNP^14^. The ANP and GC-B receptors belong to a family of GCase-coupled receptors that share a similar overall molecular configuration^21, 22^. and it is conceivable that they also share a common ligand-binding transmembrane transmission mechanism. To investigate ligand recognition and transmembrane signaling mechanisms mediated by the ANP receptor, we determined the crystal structure of the ANP receptor extracellular ANP-binding domain (ANPR) complexed with a rat partial ANP comprising amino acids 7 – 27 (rANP[7-27]) in a previous study^23^. We found that one molecule of bound ANP was flanked by two ANPR monomers. Compared with the *apo* structure^24^, ANPR caused rotation motion with no appreciable intramolecular change in either of the monomers^23^. However, low resolution of the previous datasets^23^ and the bound ANP occurs in two alternative conformations (orientations) of equal occupancy (50%) related by a two-fold symmetry, which rendered the assignment of the bound ANP inaccurate. Therefore, the ligand recognition mechanism of the receptor remained largely obscured.

This study aimed to determine the high-resolution crystal structures of the ANP receptor in complexes with rat and human full-length ANP, full-length DNP, and several ANP mutants, as well as evaluate the kinetics of the ligands used to determine the crystal structures. Our study contributes to the current knowledge on this topic in various ways and addresses gaps including ligand recognition mechanisms and transmembrane signaling mechanisms associated with the ANP receptor. The knowledge gleaned from our study can be extended to other related receptors, applied to drug discovery and design, and help further expand the research on heart failure therapies.

## Results

### Overall structures

The extracellular domain of ANPR comprising residues 1 – 435 was expressed and purified as described previously^23, 25, 26^. All structures were solved by molecular replacement using the *apo* structure (PDB accession code 1DP4^24^) as the template. Supplementary Table 1 shows crystallographic data collection and refinement statistics. Figure 1a and b shows the structures of ANPR with bound full-length human ANP (hANP[1-28]) and dendroaspis natriuretic peptide (DNP). Figure 1c and d shows their magnified views. ANP and DNP are cyclic peptides with a disulfide bond between Cys7 and Cys23 (Fig. 1e, Supplementary Fig. 1). The bound peptides are sandwiched between two monomers of ANPR with two-fold symmetry, which is similar to our previous findings of the structure in complex with rANP[7-27]^23^. The receptor monomer consists of two globular domains, a membrane distal (MD) domain and a membrane proximal (MP) domain. One Cl ion, which is indispensable for ANP binding to ANPR, is bound to the MD domain^27, 28^ and two sugar modifications are located at Asn13 and Asn395 in the MD and MP domains, respectively. The structure contains many water molecules, including the ligand-binding site (Fig. 1a, b). The structures of ANPR with bound ANP and DNP are almost identical except for one loop connecting the MD and MP domains (Supplementary Fig. 2). Other than hANP[1-28] and DNP, we determined the structures of ANPR complexed with full-length rat ANP (rANP[1-28]), truncated mutants of rANP or hANP, and deletion mutant of rANP lacking a part of the ring structure (Fig. 1e). Since two bound ligands occur in two alternative conformations (orientations) of equal occupancy (50%) related by a two-fold symmetry (Supplementary Fig. 3a, b, c), we precisely modeled the structures under the guidance of multiple difference density maps of different peptides (Supplementary Figs. 3 – 5). Details are provided in Supplementary Methods.

**Fig. 1.**
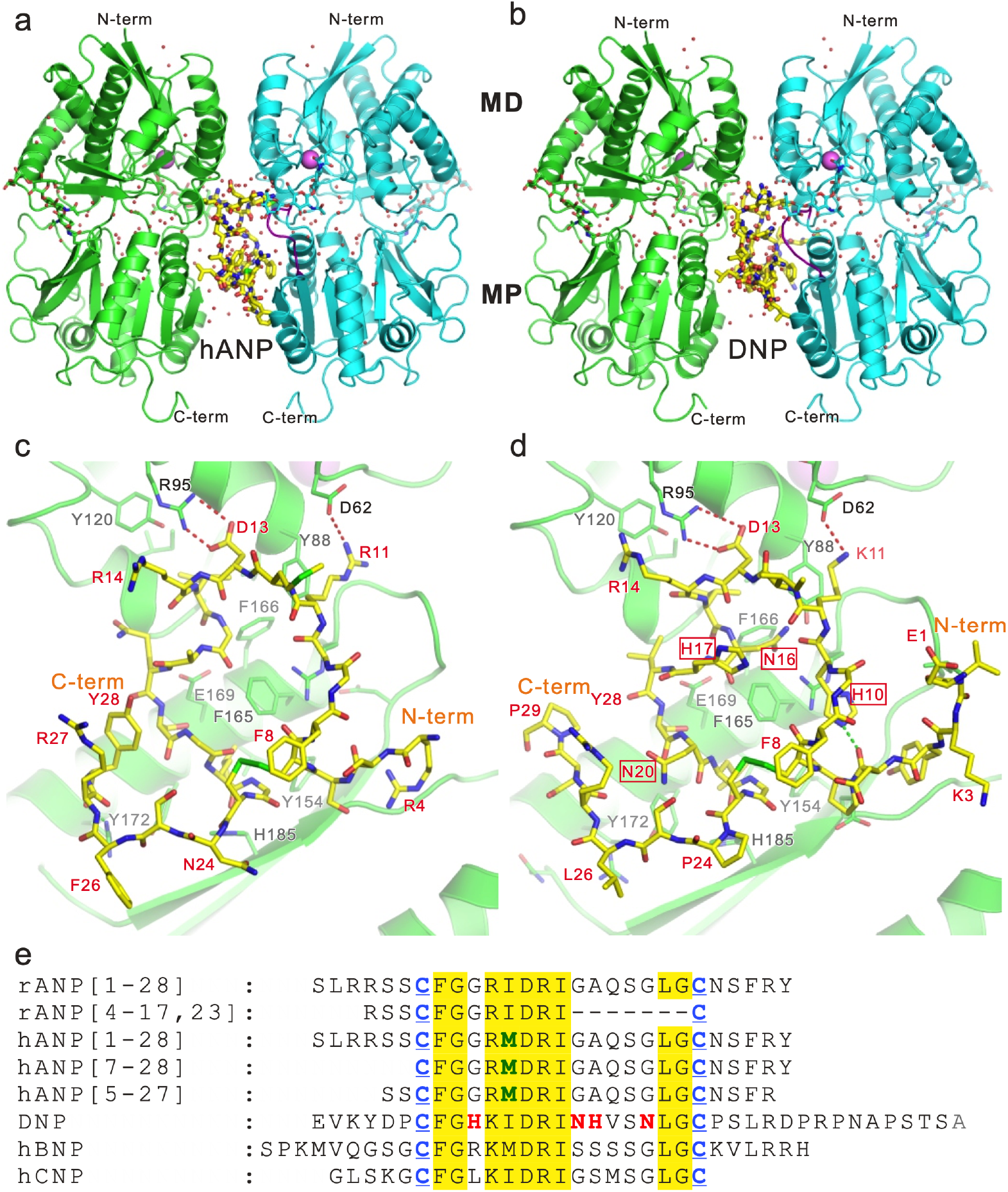
Structures of ANPR with bound natriuretic peptides. Overall structures of ANPR with bound hANP[1-28] (a) and DNP (b). ANPR is represented as a ribbon model in which monomers A and B are shown in green and cyan, respectively. Bound ligands are shown as yellow sticks. Red, oxygen; blue, nitrogen; yellow-green, sulfite atoms. Water molecules are shown as red spheres. (c, d) Magnified views around bound hANP[1-28] (c) and DNP (d), looking from the position of monomer B. (d) Amino acid residues surrounded by red squares are DNP-specific. (e) Sequence alignment of natriuretic peptides. Structures of various ANP derivatives and DNP were compared in this study with hBNP and hCNP references. Two Cys residues (blue with underscore) form an SS-bond, and the peptide forms a cyclic structure. Amino acid residues masked with yellow have conserved sequences between natriuretic peptides. Amino acid residues shown in green and red are hANP- and DNP-specific, respectively.

### Ring structure of natriuretic peptides

The ring structure of the bound ANP and DNP (amino acid residues from Cys7 to Cys23) exhibited pseudo two-fold symmetry (Fig. 2), which is not a feature of the primary sequences of natriuretic peptides (Fig. 1e, Supplementary Fig. 1). One region with two-fold symmetry was located at the top of the ring structure and the other was located at the bottom centered on the SS-bond between Cys7 and Cys23 (Fig. 2). Positively charged and nonpolar amino acid residues were positioned with pseudo two-fold symmetry centered on Asp13 at the top of the ring structures (Fig. 2a). The side chains, with positive charges in hANP[1-28], corresponded to Arg11 and Arg14, and the nonpolar region corresponded to Met12 and Ile15 (Fig. 2b). These amino acid residues in the two-fold symmetry protruded into symmetrical hydrophilic or hydrophobic pockets in the ANPR (Fig. 2b), thus stabilizing peptide binding to the receptor. Arg11 hydrogen bonds with Asp62(A), residue from monomer A of the ANPR, and the carbonyl oxygen of Asp160(A), and Arg14 hydrogen bonds with the carbonyl oxygen of Asp160(B), residue from monomer B of the ANP. Met12 and Ile15 fit into hydrophobic pockets comprising Tyr88(B), Ala111(B), Tyr120(B), Phe166(B), and Tyr88(A), Ala111(A), Tyr120(A), and Phe166(A), respectively. This scenario is similar in the ANPR with bound rANP[1-28] or DNP. Hydrophobic residues in the two-fold symmetry are the same between hANP[1-28], rANP[1-28], and DNP, except for the replacement of Met12 with Ile12 in rANP[1-28] and DNP (Fig. 2d, e, Supplementary Fig. 6a, b). Since Ile12 is nonpolar and its side chain is essentially the same size as that of Met, Ile12 easily fits into the hydrophobic pocket (Fig. 2d, e, Supplementary Fig. 6a, b). The positively charged residues in the two-fold symmetry are essentially the same between hANP[1-28], rANP[1-28], and DNP, except for the replacement of Arg11 with Lys11 in DNP (Fig. 2b, e). Another pseudo two-fold symmetry in hANP[1-28], rANP[1-28] and DNP is evident at the bottom of the ring structure (Fig 2a, d, Supplementary Fig. 6a). Nonpolar amino acid residues (Phe8 and Leu21) are positioned with pseudo two-fold symmetry centered on the SS-bond consisting of Cys7 and Cys23 (Fig. 2a, c, d, f, Supplementary Fig. 6a, c). Both nonpolar residues are firmly embedded in the hydrophobic pocket in the ANPR, thus stabilizing peptide binding to the ANPR.

**Fig. 2.**
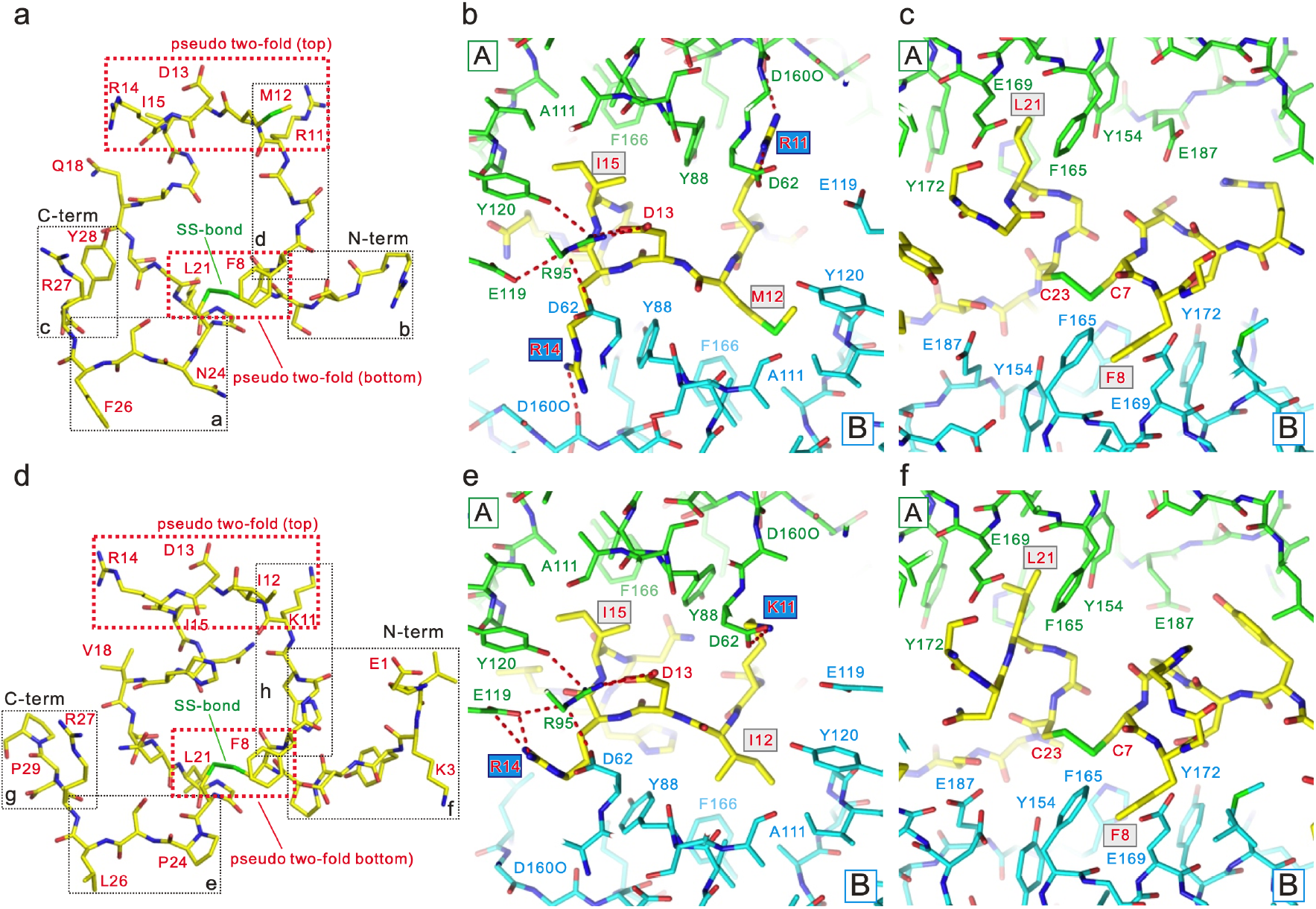
Pseudo two-fold regions of bound peptides. (a, b, c) hANP[1-28]; (d, e, f) DNP. The structure of hANP[1-28] bound to the ANPR (a) and structure of DNP (d). hANP[1-28] and DNP have pseudo-two-fold symmetry at the top and bottom of ring structures, and both regions are surrounded by red dotted boxes (a, d). The regions surrounded by black dotted boxes with alphabets a – d (a) and with alphabets a – d (d) are the regions used for the magnified views in Fig. 3. Magnified views of pseudo two-fold symmetry region at the top of the rings in hANP[1-28] (b) and DNP (d). All molecules are represented as sticks. Monomers A and B are green and cyan, respectively. The bound ligands are shown as yellow sticks. Red, oxygen; blue, nitrogen; yellow-green, sulfite atoms. Red dashed lines represent hydrogen bonds. Two independent pairs with pseudo-two-fold symmetry at the top of each ring structure are centered on Asp13. One is hydrophobic and consists of Met12 and Ile15 (hANP[1-28]) or Ile12 and Ile15 (DNP), the other is hydrophilic and consists of Arg11 and Arg14 (hANP[1-28]) or Lys11 and Arg14 (DNP). Both pairs can enter symmetrical hydrophobic or hydrophilic pockets of ANPR monomers. Magnified views of pseudo two-fold symmetry region at the bottom of rings in hANP[1-28] (c) and DNP (e). Hydrophobic pairs of Phe8 and Leu21 centered on the disulfide bond composed of Cys7 and Cys23 have pseudo two-fold symmetry. Hydrophobic pairs can enter symmetrical hydrophobic pocket of ANPR monomers.

### Architecture of the ligand recognition by ANPR

Since the structures of the bound hANP[1-28] and rANP[1-28] are essentially identical except for amino acid residue 12, we used the structure of hANP[1-28] as the standard for comparisons with DNP. Four common features are essential for ANP and DNP binding to the ANPR. One is the pseudo two-fold symmetry at the top of the ring structure, consisting of positively charged residues and nonpolar residues centered on Asp13 (Fig. 2a, b, d, e, Supplementary Fig. 6a, b). The second is the pseudo two-fold symmetry at the bottom of the ring structures consisting of Phe8 and Leu21 centered on the SS-bond (Fig. 2a, c, d, f, Supplementary Fig. 6a, c). The third is Asp13 at the center of the pseudo two-fold symmetry at the top of the ring structure, at the boundary between monomers A and B of the ANPR. There, it forms a tight salt bridge with Arg95(A), which is a key step for dimer formation. In fact, Arg95(A), a salt bridge partner of D13, acts as a mediator that binds to not only sidechains from monomer A (Glu119 (A) and Tyr120 (A)) but also those from monomer B (Asp62 (B)) (Fig. 1c, d, Fig. 2b, e). The fourth feature is a pseudo β-sheet between 3 amino acid residues immediately after the SS-bond of the ring structure (amino acid residues 24 – 26) and monomer B of the ANPR (Fig. 3a and e). The essential feature of this pseudo β-sheet was shown to be common to both ANP and DNP. The carbonyl oxygen atom of Asn24(ANP) or Pro24(DNP) was shown to form hydrogen bonds with the nitrogen atoms of Glu187(B), while Phe26(ANP) or Leu26(DNP) forms hydrogen bonds with the carbonyl oxygen atom of Glu187(B) (Fig. 3a and e). Moreover, ANP and DNP were shown to have additional hydrogen bonds to reinforce the pseudo β-sheet. However, the residues used for hydrogen bonding in ANP and DNP slightly differed due to the difference in the amino acid residues between them. Asn24 and Ser25 formed hydrogen bonds with the carbonyl oxygen of His185(B) and with Glu187(B), respectively, in ANP, whereas Ser25 and the carbonyl oxygen of Leu26 formed hydrogen bonds with Glu187(B) and His195(B), respectively, in DNP.

**Fig. 3.**
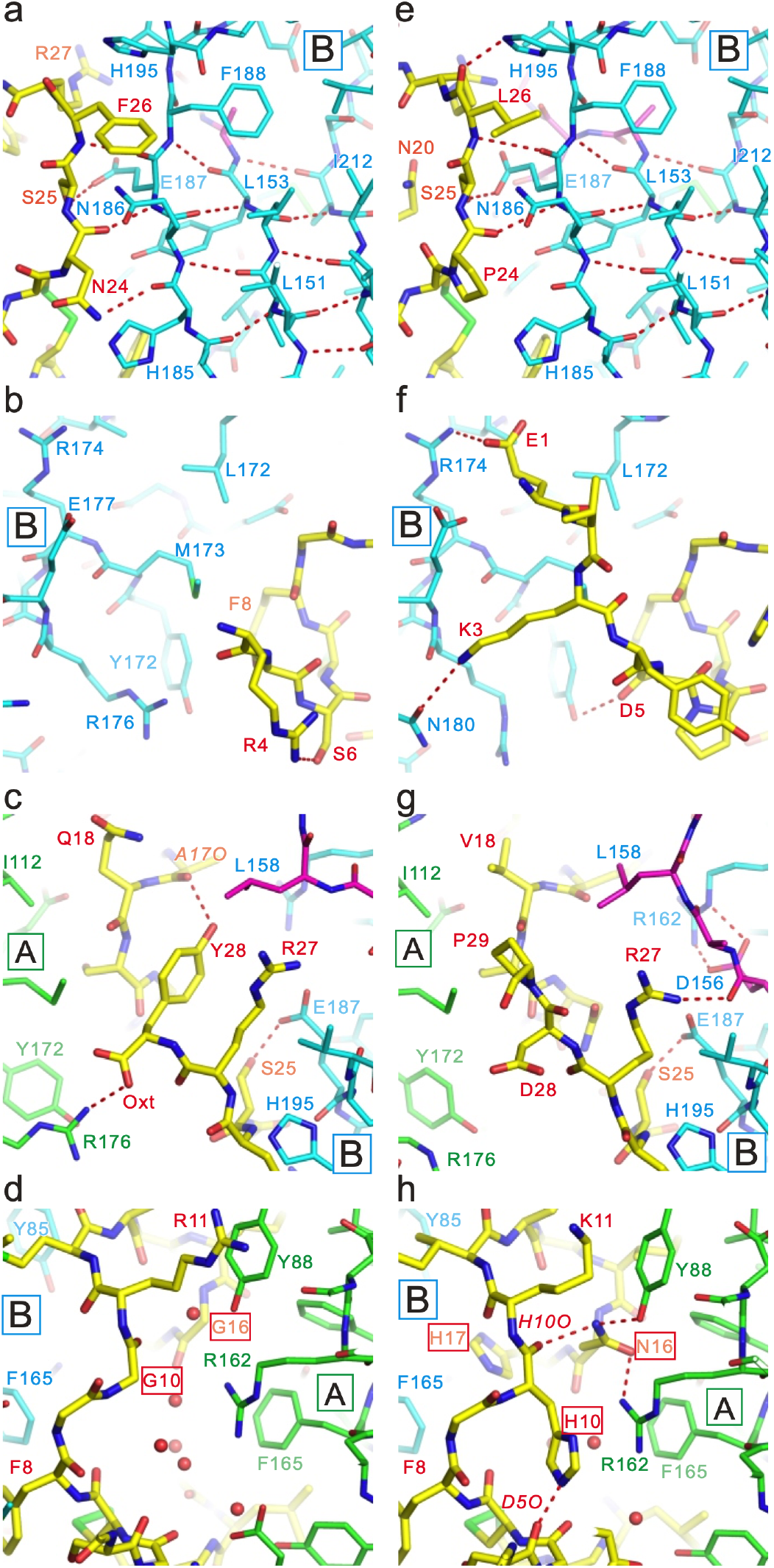
Details of the binding of bound peptides. (a, b, c, d) Bound hANP[1-28]. (e, f, g, h) Bound DNP. Region of pseudo β-sheet between ANPR and hANP[1-28] (a) or DNP (e). N-terminus region of hANP[1-28] (b) or DNP (f) before SS-bond. C-terminus region after pseudo β-sheet of hANP[1-28] (c) or DNP (g). Amino acid residues around DNP-specific region in hANP[1-28] (d) and DNP (h).

The N-terminus sequences of ANP and DNP and the directions of the primary chain in each peptide are quite different (Fig. 3b and f). The N-terminus of DNP is well resolved compared with ANP, and three hydrogen bonds are located among Glu1(DNP)- Arg174(B), Lys3(DNP)-Asn180(B), and Asp5(DNP)-Tyr172(B) (Fig. 3f). Except for the first three residues, those in ANP were well resolved. Arg4 formed a hydrogen bond with Ser6, but not with ANPR (Fig. 3b). These findings indicate that the N-terminus of ANP is more flexible than that of DNP. In contrast, the C-termini of ANP and DNP play important roles in binding to ANPR, although the sequences of these termini largely differed (Fig. 1e). The last residue in ANP, Tyr28, plays a key role in the formation of the ANPR/ANP complex. The carboxyl terminal oxygen atom of Tyr28 formed a hydrogen bond with Arg176(A), which belongs to the other monomer of the ANPR (Fig. 3c). This hydrogen bond is important for the formation of tight ANPR/ANP complexes. In addition, the side chain of Tyr28 formed a hydrogen bond with the carbonyl oxygen atom of Ala17 and helps to stabilize the ring structure of ANP. Induced-fit to DNP occurs in ANPR, and loop 155 – 161 of monomer B comes close to Arg27(DNP) (Fig. 3g, Supplementary Fig. 2b). As a result, Asp156(B) in the loop can form a hydrogen bond with Arg162(B), and Arg27(DNP) can form a hydrogen bond with carbonyl oxygen atom of Asp156(B) (Fig. 3g). No specific bond exists between ANPR and Asp28(DNP) or Pro29(DNP), and amino acid residues after 30 were not well resolved in DNP. The most characteristic difference between ANP and DNP was seen in the four DNP-specific amino acid residues (His10, Asn16, His17 and Asn20) within the ring structure of the peptides (Fig. 3h), wherein the corresponding residues in ANP were all Gly or Ala (Gly10, Gly16, Ala17 and Gly20) (Fig. 3d). His10(DNP) formed a hydrogen bond with the carbonyl oxygen of Asp5(DNP), and the direction of the His-ring of His10(DNP) was fixed in DNP (Fig 3h). As a result, His10(DNP) constituted a stack of π interactions between Arg162(A) and Phe165(A). Asn16(DNP) formed a hydrogen bond between the carbonyl oxygen atoms of His10(DNP), Tyr88(A) and Arg162(A) (Fig. 3h). Although water molecules surrounded the corresponding residues Gly10 and Gly16 in ANP, no specific hydrogen bond between ANP and the ANPR was evident around these residues (Fig. 3d). It is reported that the affinity of DNP for the ANPR is stronger than that of ANP^29^. Our findings supported that DNP has higher affinity to the ANPR than ANP.

### Identification of key residues in ANP involved in ANPR binding

We further investigated the details of substrate binding to the ANPR using blue native polyacrylamide gel electrophoresis (BN-PAGE). We detected a band for ANPR without a ligand in the region of ANPR monomer, but the band for the ANPR complexed with physiological ligands (hANP[1-28], rANP[1-28], rBNP, or DNP) resolved at the region of the dimer in ANPR (Fig. 4a). In contrast, when CNP, which is not a ligand for this receptor, was added to ANPR, the band for the ANPR was detected as a monomer (Fig. 4a). We applied BN-PAGE to the various ANP mutants and investigated how the locations of the mutations affect ANPR dimerization. Focusing on the N-terminus deletion mutants of ANP, the findings of rANP[3-28] and rANP[1-28] were almost identical. In contrast, dimerization was induced in hANPR[5-28] and hANP[7-28], but the bands were tailed slightly to the region of the ANPR monomer (Fig. 4a). We showed that Arg4(ANP) formed a hydrogen bond with Ser6(ANP) in ANPR complexed with hANP[1-28] (Fig. 3b) and deleting residues 4 – 6 in ANP led to structural instability of bound ANP, which could affect ANPR dimerization.

**Fig. 4.**
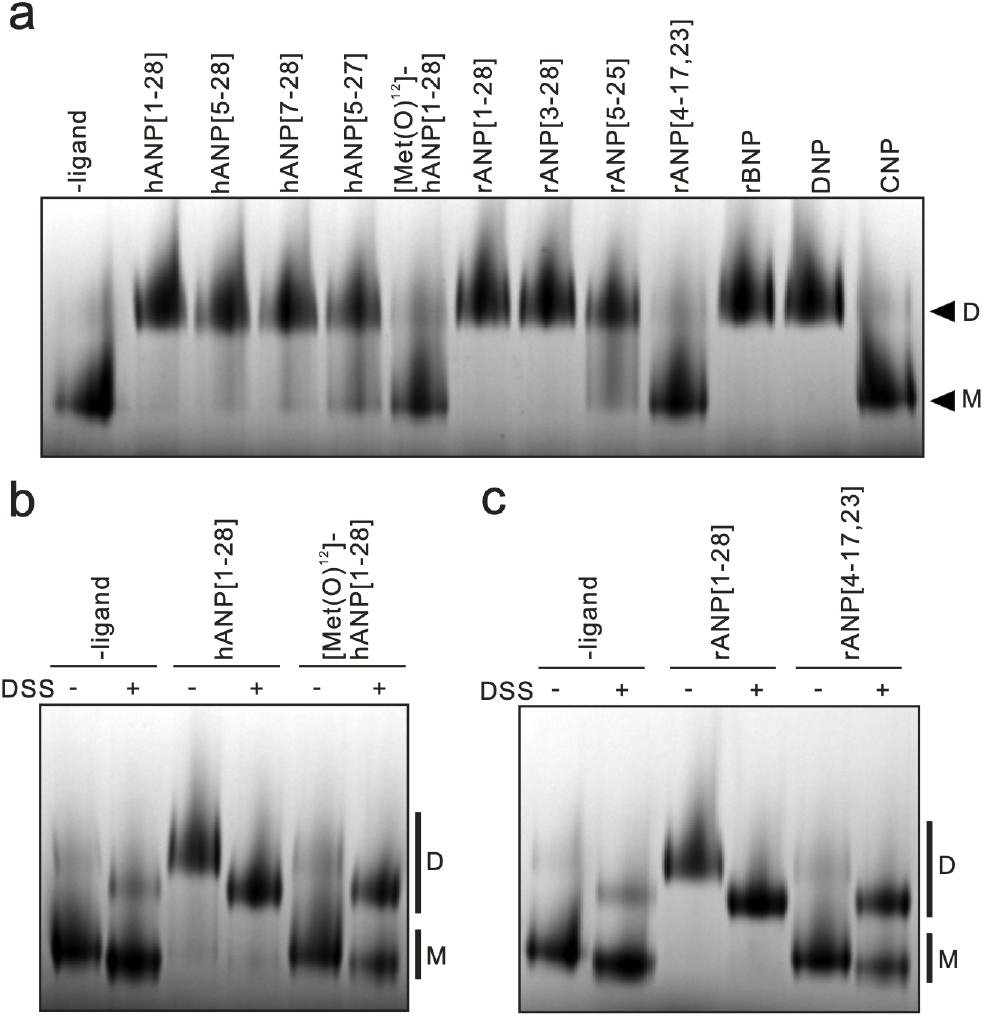
Ligand-dependent dimerization of ANPR. (a) Following the incubation of ANPR with various ligands, complexes were resolved by BN-PAGE and stained with Coomassie brilliant blue (CBB). M, position of ANPR monomer; D, position of ANPR dimer. (b, c) Effects of the cross-link with/without disuccinimidyl substrate (DSS) prior to BN-PAGE. Following incubation of ANPR with various ligands, complexes were cross-linked with DSS, then resolved by BN-PAGE and stained with CBB. (b) Results with Met(O)^12^-hANP[1-28] and controls without ligand or hANP[1-28]. (c) Results with rANP[4-17, 23] and controls without ligand and rANP[1-28].

Mutant hANP[5-27] with a deleted C-terminus induced dimerization, but several bands were shown to be tailed to the region of the ANPR monomer compared with hANP[5-28]. (Fig. 3c). Although most of the pseudo β-sheet forming between the ANPR is missing from rANP[5-25], the electrophoretic profile of rANP[5-25] was almost identical to that of hANP[5-27] (Fig. 4a).

Met(O)^12^-hANP[1-28] comprises a methionine sulfoxide molecule formed through sulfur oxidation in Met12 of hANP[1-28], which inhibits its ANP receptor agonistic properties^30, 31^. The crystal structure complexed with hANP[1-28] (Fig. 2a, b) showed that the Met12(hANP) and Ile15(hANP) pair constituted a pseudo-two-fold symmetry as hydrophobic residues at the top of the ring structure. Since methionine sulfoxide is hydrophilic, the two-fold symmetry as a pair of hydrophobic amino acid residues at this location breaks, and the resultant hydrophilic side chain would not fit the hydrophobic pocket (Fig. 2b). Therefore, dimerization was less likely to occur (Fig. 4a). However, unlike the ANPR without ligand, the band was shown to have tailed slightly to the region of the ANPR dimer (Fig. 4a), since Met(O)^12^-hANP[1-28] still possesses another pseudo-two-fold symmetry at the bottom of the ring structure. rANP[4-17, 23] lacks the important hydrophobic Leu21 residue that contributes to the pseudo two-fold symmetry with Phe8 at the bottom of the ring structure, and is known for its inability to behave as an ANP receptor agonist^32^. Therefore, dimerization was less likely to have occurred, but the mobility shifts of ANPR with rANP[4-17, 23] and Met(O)^12^-hANP[1-28] were almost identical (Fig. 4a), since rANP[4-17, 23] kept another pseudo-two-fold symmetry at the top of the ring structure.

The mobility shifts in ANPR with Met(O)^12^-hANP[1-28] and rANP[4-17, 23] completely differed from those in ANPR without a ligand. Therefore, we cross-linked the ANPR with disuccinimidyl substrate (DSS), having spacer arm length of 11.4 Å, before applying BN-PAGE to clarify these phenomena. According to the crystal structure, the distance between Nζ-Lys206(A) and Nζ-Lys206(B) of the ANPR became ~ 11 Å upon ligand binding (Fig. 1a and b). Figure 4b and c shows the results of BN-PAGE analysis of ANPR with and without DSS cross-linking. The positive controls were ANPR with hANP[1-28] (Fig 4b) or rANP[1-28] (Fig. 4c), and the negative control was ANPR without ligand (Fig. 4b, c). The main band of ANPR without the ligand resolved at the monomer region, regardless of the presence or absence of DSS, although the band of ANPR modified with DSS shifted slightly (Fig. 4b, c). In contrast, the main bands of ANPR with rANP[1-28] or hANP[1-28], resolved at the dimer region regardless of the presence or absence of DSS (Fig. 4b, c). Like ANP without the ligand modified with DSS, the band of ANPR modified with DSS shifted (Fig. 4b, c). However, bands for the dimer, which never appeared in ANPR without a ligand even after modification with DSS, were clearly visible for ANPR with Met(O)^12^-hANP[1-28] or rANP[4-17, 23] (Fig. 4b and c). These results indicate that although Met(O)^12^-hANP[1-28] or rANP[4-17, 23] can bind to the ANPR, the main band of the ANPR appeared to resolve in the region of the monomer and tailed slightly to the region for the dimer of the ANPR on BN-PAGE because of its low affinity.

### ANP with lost two-fold symmetry

We identified one pseudo two-fold symmetry each at the top and bottom of the ring structure of natriuretic peptides and determined their roles. Based on the crystal structures and the results from the BN-PAGE analysis, the symmetry at the top appeared to be key for attracting monomers A and B of ANPR, whereas the role of the bottom remained obscured. To investigate this, we aimed to determine the crystal structure of ANPR complexed with rANP[4-17, 23] since it was shown that rANP[4-17, 23] can bind to the ANPR (Fig. 4c). The unit cell dimension of the crystal of ANPR complexed with rANP[4-17, 23] notably differed from that of any other crystal. In fact, the *c*-axis was 9 Å longer than that of typical ligand complexes (Supplementary Table 1). Although the resolution was relatively low (3.2 Å), we obtained a clear electron density map for the bound rANP[4-17, 23] (Supplementary Fig. 7). The structure was significantly smaller for bound rANP[4-17, 23] than for rANP (Fig. 5a, Supplementary Fig. 7b and c). Figure 5b shows the location of monomer B of ANPR in the *apo* state, with bound rANP[1-28] and bound rANP[4-17, 23] fitted between monomer A of the ANPR. Monomer B of rANP[1-28] and rANP[4-17, 23] rotated around the pivot point with respect to the *apo* structure (Fig. 5b; red circle). Monomer B of the ANPR complexed with rANP[1-28] rotated 22° relative to the ANPR without the ligand. Monomer B of the ANPR complexed with rANP[4-17, 23] rotated an extra 3° compared with the ANPR complexed with rANP[1-28] (Fig. 5b). The pseudo two-fold symmetry located at the top of the ring structure of the rANP[4-17, 23] was maintained with slight distortion (Fig. 5c). In the pseudo two-fold symmetry at the bottom of the ring centered on the SS-bond (Phe8 and Leu21 pair), Phe8 was embedded in the hydrophobic pocket of monomer A (Fig. 5d). However, rANP[4-17, 23] lacks Leu21, thus no residues were embedded in the hydrophobic pocket of monomer B. These findings indicate that Leu21 in rANP[1-28] functions to stop the rotation of monomer B at 22° by embedding it in the hydrophobic pocket. Because rANP[4-17, 23] lacks Leu21, it also lacks such a brake, which resulted in an extra 3° of rotation compared with rANP[1-28] (Fig. 5d). In conclusion, pseudo two-fold symmetry at the bottom region of the ring structures of ANP or DNP, comprising Phe8 and Leu21 centered on the SS-bond, could act to stop rotation at a specific angle.

**Fig. 5.**
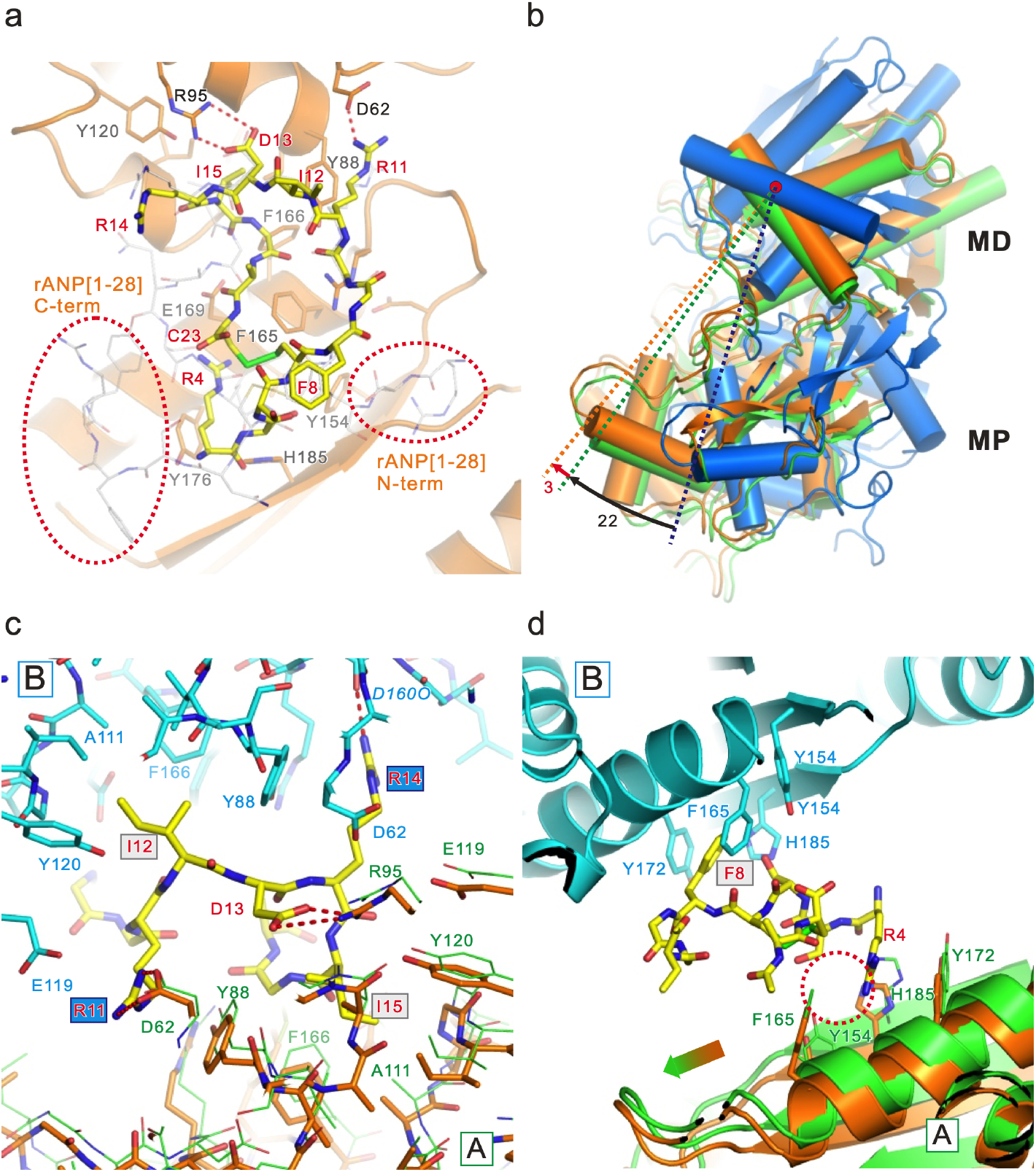
Crystal structure of ANPR with bound ligand lacking two-fold symmetry. (a) Magnified views of the ANPR structure with bound rANP[4-17, 23] looking from monomer B. The ANPR is shown as a ribbon model. The bound rANP[4-17, 23] is shown as a yellow stick. rANP[1-28] bound to ANPR is overlaid as a white stick. Red, oxygen; blue, nitrogen; yellow-green, sulfite atoms. (b) Overlay of ANPR in *apo* (blue), with bound rANP[1-28] (green) and bound rANP[4-17, 23] (orange). These three structures were fitted using monomer A of the ANPR. The ANPR with bound rANP[1-28] rotates 22° around the red dot with respect to *apo* structure. The ANPR rotates 3° more with bound rANP[4-17, 23] than with bound rANP[1-28]. (c) Magnified view of pseudo two-fold symmetry region at the top of the ring in rANP[4-17, 23]. Monomers A and B are orange and cyan, respectively. Yellow sticks, bound rANP[4-17, 23]. Red, oxygen; blue, nitrogen; yellow-green, sulfite atoms. Thin green stick, monomer A with bound rANP[1-28]. (d) Magnified view of pseudo two-fold symmetry region at the bottom of the ring in rANP[4-17, 23]. Ribbon model shows ANPR. Orange and cyan, monomers A and B, respectively. Yellow sticks, bound rANP[4-17, 23]. Red dotted circle, location corresponding to absent pseudo two-fold symmetry of bound rANP[4-17, 23]. No side chain exists to fit inside hydrophobic pocket on monomer A of ANPR. Therefore, ANPR rotates more (arrow) upon binding to rANP[4-17, 23].

## Discussion

Using high-resolution structures of ANPR with various bound natriuretic peptides having many water molecules, we elucidated the substrate recognition and signal transmission mechanism of ANPR. Accurate modeling of the bound peptides was the most challenging, since two bound ligands attach in two alternative conformations (orientations) of equal probability (50%) related by a two-fold symmetry (Supplementary Fig. 3a, b, c). Peptides, including the physiological ligand ANP and snake venom DNP, when bound to ANPR notably had a similar pseudo-two-fold symmetry and closely identical overall structures. The critical role of amino acid residues comprising the pseudo-two fold symmetry is consistent with the finding that these residues are essential for the hormonal activity of ANP^33^. The structure of BNP, a physiological ligand of ANPR, complexed with ANPR remains unknown. However, considering its sequence identity with ANP or DNP (Fig. 1e, Supplementary Fig. 1), BNP could bind to ANPR with pseudo two-fold symmetry, in the same manner as ANP. The physiological ligand for the B-type natriuretic receptor (GC-B receptor), CNP, which plays a critical role in cartilage formation^16^, also has ~ 65% of sequence identity with ANP and BNP in the ring structure (Fig. 1e, Supplementary Fig. 1). Although the structure of CNP complexed with the GC-B receptor remains unknown, CNP might also bind to the GC-B receptor with pseudo-two-fold symmetry, in the same manner as ANP. Guanylyl cyclase-C (GC-C) receptor belongs to a family of GCase receptors that is thought to have the same transmembrane signaling mechanism as the ANPR^34^. The physiological ligands for the GC-C receptor are guanylin^35^ and uroganylin^36^, which both have two disulfide bonds. The GC-C receptor is also known as the receptor for a heat-stable enterotoxin that has three disulfide bonds^34, 37^. Although the structure of the GC-C receptor with bound ligands remains unknown, the ligands might bind with pseudo two-fold symmetry, in the same manner as ANP.

The potent natriuretic and diuretic peptide, DNP, is similar to ANP and BNP and induces an increase in urinary and plasma cGMP levels. Compared with ANP, BNP, and CNP, DNP has elongated N- and C-termini and is more stable against neutral endopeptidase^38^, which is the primary inactivator of natriuretic peptides. Based on the enhanced stability and high potency of DNP, chimeras of DNP and other natriuretic peptides have been created and assessed in clinical trials for heart failure^39, 40^. Thus, understanding the difference between the binding of ANP and DNP to ANPR is essential to future drug discovery and research for heart failure therapies. The key difference between ANP and DNP lies in the four DNP-specific amino acid residues (His10, Asn16, His17 and Asn20) within the ring structure (Fig. 1e). Since the corresponding residues in ANP were all small Gly or Ala, and most are also conserved in BNP and CNP (Fig. 1e), these small residues were considered essential for increasing the flexibility of the main peptide chain. However, a comparison of ring structures between ANP and DNP did not reveal any appreciable difference between them (Fig. 2a, b). This observation suggests that His10, Asn16, His17, and Asn20 are DNP-specific and play important roles in binding to the ANPR. A notably sufficient space between these corresponding residues and ANPR was filled with water molecules (Fig. 3h). In contrast, the space between DNP and ANPR was occupied by substituted His and Asn without steric hindrance, and water molecules in ANP binding were completely excluded. Moreover, the substituted His and Asn form hydrogen bonds with ANPR or DNP themselves, suggesting that they play important roles in tightly binding to ANPR or in reinforcing the structure of DNP. Thus, these findings suggest that DNP can bind more tightly than ANP to the ANPR. In fact, reports of concentration-dependent GCase activity generated by ANP or DNP in cultured cells showed that the EC_50_ was 10-fold higher for DNP than for ANP^41^. Due to enhanced stability and affinity^29, 38^, DNP binding could be more stable and longer-lasting than that of ANP. Thus, higher production of cGMP was induced by DNP rather than by ANP. This could potentially explain the functioning of DNP as a snake venom.

The extracellular ligand-binding domain of C-type natriuretic peptide receptor (NPRC), also known as natriuretic peptide clearance receptor, shows ~30% similarity to the extracellular ligand domain of the ANP receptor. Unlike the ANP receptor, NPRC lacks the GCase domain and is not coupled to GCase^42^. The structures of NPRC in *apo*, with bound ANP, BNP, or CNP have been previously determined^43, 44^. The structure of the NPRC monomer consists of MD and MP domains that are similar to ANPR. However, the structural changes upon ligand binding are completely different. Whereas ANPR causes rotation motion without any structural changes in the monomer upon ligand binding^23^, NPRC upon ligand binding causes a pinching motion of the MP; that is, the MP domain swings onto the bound ligand, leaving the MD domain and its dimerized structure essentially unchanged^43^. A plausible explanation for the difference in structural change between the two receptors following ligand binding could be associated with their physical functions. In fact, the NPRC binds ANP, BNP, and CNP as well as some of their biologically inactive fragments with equally high affinity and removes excess peptides from the blood circulation^32, 45, 46, 47^. The pinching motion in the NPRC might facilitate the acceptance of a wider range of ligands than the rotation in the ANPR. Our structural comparison of the bound ligands between NPRC and ANPR showed that peptides bound to NPRC do not have the pseudo two-fold symmetry that we identified in ANPR (Supplementary Fig. 8). In fact, NPRC has been shown to bind fragments of the natriuretic peptide, and some of these peptides lack the elements that can form pseudo two-fold symmetry in the peptides^31, 32^. To bind to peptides lacking pseudo two-fold symmetry and to transmit its binding signal into the cell, the NPRC might require a signal transmission mechanism that does not depend on ligands having a pseudo two-fold symmetry.

Although acute heart failure can be treated with hANP[1-28], there are disadvantages associated with this treatment approach. Continuous infusion for longer periods is necessary because the half-life of hANP[1-28] in the bloodstream is only 0.5 – 4 minutes^48, 49^. Thus, the development of compounds with a longer half-life is necessary. Moreover, hANP[1-28] can only be dissolved in water for injection and not with saline. Thus, compounds that are soluble in saline are also desired. A comparison of the structures of ANPR with bound ANP and DNP provides important clues to new drug development for heart failure. Moreover, since ANPR forms a homodimer, two ANPR facing each other naturally have a two-fold symmetry. Therefore, if a compound with perfect two-fold symmetry can cause rotation of ANPR, it could become the foundation for a drug to treat heart failure. However, the requirements for activation of ANPR not only includes rotation *per se*, but also the correct angle of the rotation. This is considered one of the primary reasons why ANPR-activating drugs are not currently commercially available. Moreover, the results of the DNP binding showed that ligands with higher affinity than that of ANP for ANPR lead to excessive cGMP production, which can be toxic. Thus, compounds with moderate affinity and two-fold symmetry to rotate the ANPR dimer to an appropriate angle could be suitable candidates for new drugs. Therefore, our structures can be relied upon for the development of new drugs for heart failure therapy.

Soluble GCase (sGC) has ~ 51% sequence identity with the GCase domain of the ANP receptor. Structures of full-length sGC in both inactive and active states were recently solved using cryo-electron microscopy^50^. sGC is a heterodimer composed of two subunits (α and β), that are each composed of four domains: an N-terminal heme nitric oxide/oxygen (H-NOX) domain, a Per/Arnt/Sim (PAS)-like domain, a coiled-coil (CC) domain, and a catalytic (CAT) domain holding GCase activity. The binding of ligands to the sGC induced large conformational change of G-NOX and PAS domains. Repositioning of the domains lead to a straightening of the CC domains, which, in turn, use the motion to move the CAT domains into an active conformation. The mechanism by which the repositioning at the ligand binding site made GCase active through the structural change in the CC domain in sGC provides insight into the mechanism wherein the extracellular ligand binding in the ANP receptor activates intracellular GCase domain through a single transmembrane helix.

Figure 6 shows a schema of a hypothetical signaling mechanism generated by the ANP receptor. Intracellular GCase domains (GCD) form a dimer, but a gap exists between the GCD facing each other in the *apo* state. Because of this gap, the receptor is inactive in the *apo* state (Fig. 6a). Upon ligand binding, ANPR causes each extracellular ligand binding domain to rotate 11°, and this motion is transmitted to the intracellular domain through the transmembrane helix. This results in a change of the face between the dimer GCD, and GCase activation (Fig. 6b). Ligands bind to the ANPR with pseudo-two-fold symmetry at the top and bottom regions of the ring structure. Two-fold symmetry at the top of the ring consists of two pairs of side-chains with different properties (hydrophobic, Met12(Ile12)/Ile15 and hydrophilic, Arg11/Arg14) centered on Asp13. This symmetry at the top and Asp13 are considered to largely contribute to ANPR dimerization (Fig. 6b). In contrast, rANP[4-17, 23] lacks pseudo two-fold symmetry at the bottom of the ring structure (Fig. 6c). Because rANP[4-17, 23] retains the pseudo two-fold symmetry at the top of the ring structure, rANP[4-17, 23] could bind to the ANPR (Fig. 6c), albeit relatively weakly (Fig. 4), and was observed to be relatively distorted compared with that of ANP or DNP binding (Fig. 5). Two-fold symmetry at the bottom of the ring comprises a hydrophobic pair consisting of Phe8 and Leu21 (Fig. 6b). Both residues fitted into the non-polar pockets, leading to precise control of the 11° rotation of each monomer of the ANPR (Fig. 6b). Because rANP[4-17, 23] does not have pseudo-two-fold symmetry at the bottom, Phe8 can fit into one non-polar pocket, but the other cannot be filled. As a result, the rotation could not be stopped at the correct angle, so it continued for ~ 3° (Fig. 6c). These findings suggest that lower pseudo two-fold symmetry functions in decelerating the rotation of the ANPR (Fig. 6c).

**Fig. 6.**
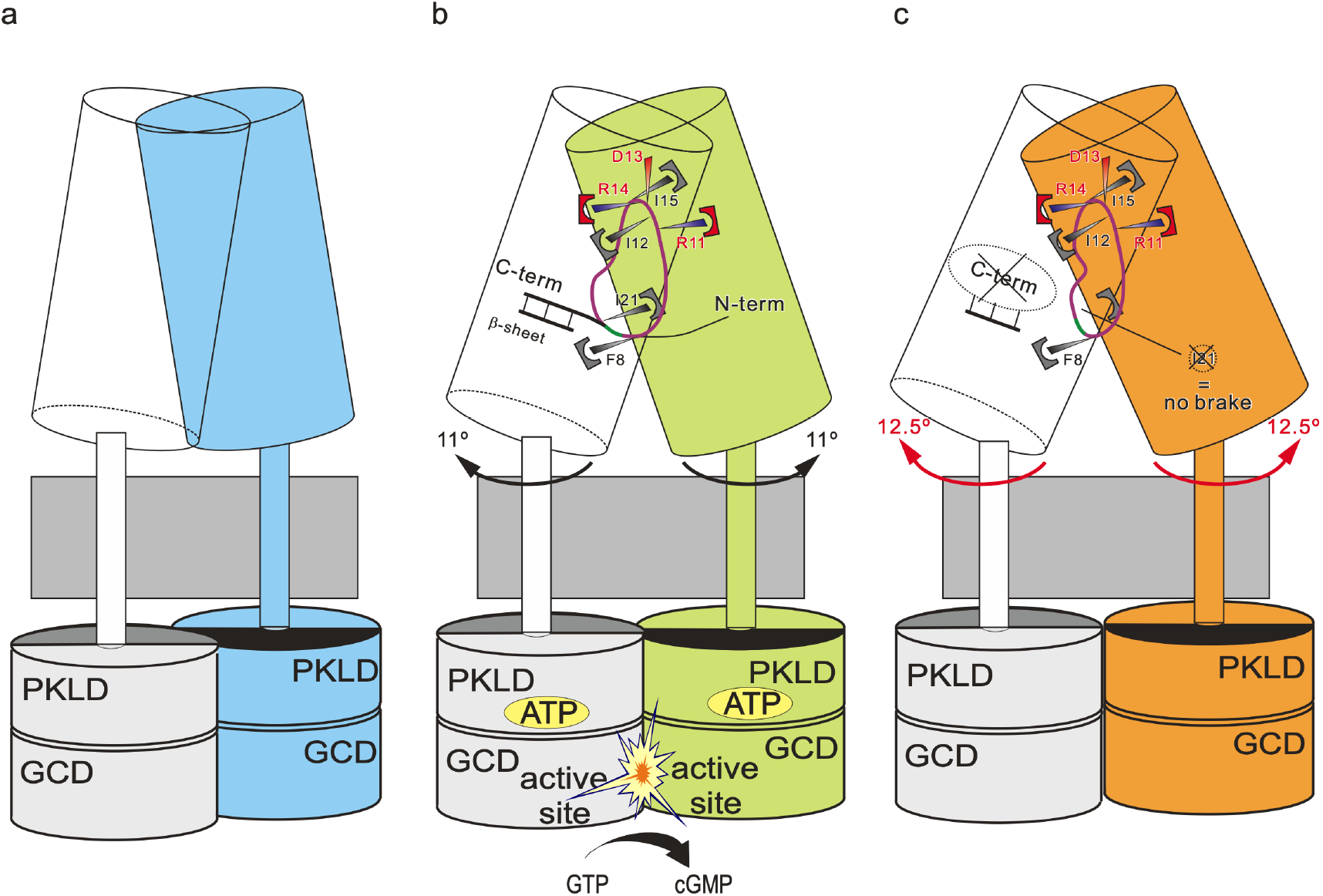
Hypothetical schema of signaling mechanism generated by ANPR. (a) Full-length ANP receptor in *apo* state, (b) with bound rANP[1-28] and (c) with bound rANP[4-17, 23].

We determined the structures of ANPR complexed with various ligands. The bound peptides possessed pseudo two-fold symmetry at two regions (top and bottom) in their ring structures. They played key roles in both the tight binding of the ligands to the ANPR as well as the signal transmission mediated by the ANPR. Kinetic and comparison studies of ANP and DNP binding to the ANPR deepen our understanding of the ligand recognition mechanism mediated by ANPR, which, in turn, may facilitate the development of novel treatment options for heart failure.

## Methods

### Production and purification of the ANP receptor

Stable cell lines expressing high levels of the ANPR were established as previously described^26^. cDNA encoding rat ANP receptor (GenBank ID: NM_012613) was purchased from OriGene Technologies Inc. (Rockville, MD, USA). Sequences, including signal sequences encoding the extracellular hormone binding domain of the ANP receptor (ANPR, amino acid residues, 1 – 463) were amplified by PCR using the sense and antisense oligonucleotide primers (underlines: restriction sites) (5′ - GGATCC<u>CTCGAG</u>CTAGTCTTGGTTGCAGGC-3′ and 5′ - TGCTCC<u>ACCGG</u>TGTCTTGGTTGCAGGCTG-3′). Each PCR product was subcloned into the NheI and XhoI sites of a cytomegalovirus promoter-driven pIRES2-AcGFP1 vector (Clontech Laboratories Inc., Mountain View, CA, USA). HEK293T cells were co-transfected with the construct and pPUR (Clontech) using Lipofectamine 2000 (Invitrogen, Carlsbad, CA, USA). The transfected cells were selected in growth medium containing puromycin (0.2 μg/mL) and cultured for two weeks. The top 10% of the cell population with the most intense GFP fluorescence was selected using a FACS Vantage SE cell sorter (Becton Dickinson and Co., Franklin Lakes, NJ, USA). The selected cells were cultured with puromycin (0.2 μg/mL) for 1 – 2 weeks then another selection cycle proceeded. Thereafter, ANPR was affinity purified as previously described^23, 25^.

### Crystallization, data collection, and structure determination

hANP[1-28], hANP[5-28], hANP[7-28], hANP[5-27], Met(O)^12^-hANP[1-28], rANP[1-28], rANP[3-28], rANP[5-25], hBNP and hCNP were purchased from the Peptide Institute (Osaka, Japan). We purchased DNP from Phoenix Pharmaceuticals Inc. (Burlingame, CA, USA) and rANP[4-17, 23] from American Peptide Co. (Sunnyvale, CA, USA). hANP[1-28], hANP[7-28], hANP[5-27]. rANP[1-28], DNP and rANP[4-17, 23] were used for the crystallization. The ANPR complex (10 – 15 mg/mL) and natriuretic peptides were crystallized by hanging drop vapor diffusion at room temperature with 1.2 – 1.6 M sodium malonate in 0.1 M MES buffer, pH 6.3 – 6.6. Crystals were quickly soaked in high concentrations of sodium malonate and frozen in liquid nitrogen.

Diffraction data at a wavelength of 0.9 Å were collected from crystals cooled to 100 K at BL41XU of SPring-8 using a ADSC Quantum 315 or Rayonix MX225HE CCD (charge-coupled device) detector. All data were processed using HKL2000^51^. Diffraction data from the two crystals with high resolution were averaged for each dataset. Crystal structures were determined by molecular replacement using *apo* ANPR (PDB ID: 1DP4) using Molrep^52^. Models were built manually using Coot, then atomic models were refined using CNS^53^ followed by Phenix^54^. Fourier maps were calculated using the CNS^53^. Structure figures were prepared using PyMol 2.4.0a (The PyMOL Molecular Graphics System, http://www.pymol.org).

### Blue native polyacrylamide gel electrophoresis (BN-PAGE) analysis

ANPR (80 pmol) was incubated with 40 pmol of peptide in 10 μl of 1X NativePAGE Sample Buffer (Invitrogen) for 30 minutes at 25 °C. Samples were applied on NativePAGE 4% – 16% Bis-Tris Protein Gels (Invitrogen) and resolved using 1X NativePAGE Anode Buffer (Invitrogen) and 1X NativePAGE Dark Blue Cathode Buffer containing Coomassie G-250 (Invitrogen), as per manufacturer instructions. For assays involving a cross-linker, 8 nmol of disuccinimidyl suberate (DSS) (Thermo Fisher Scientific Inc., Waltham, MA, USA) was added to the reaction mixture prior to incubation.

## Data availability

The coordinates and structure factors for ANPR complexed with ligands have been deposited in the Protein Data Bank (PDB) under the accession codes 7BRG for rANP[1-28] complex, 7BRH for hANP[1-28] complex, 7BRI for DNP, 7BRJ for hANP[7-28], 7BRK for hANP[5-27], and 7BRL for rANP[4-17,23] data.

## Acknowledgements

We thank Dr. Chikashi Toyoshima (University of Tokyo) for supporting the initial setup of this investigation, and Mariko Kurakata (University of Tokyo) for technical assistance. We thank K. Hasegawa and H. Okumura of the Japan Synchrotron Radiation Research Institute (JASRI) for their help in collecting diffraction data at BL41XU of SPring-8. This study was supported by JSPS KAKENHI Grant No. JP22570110 to H.O.; Platform Project for Supporting Drug Discovery and Life Science Research (Basis for Supporting Innovative Drug Discovery and Life Science Research (BINDS) (Grant Nos. JP17am0101080, JP18am0101080 and JP19am0101080) to H.O., and a research grant from the Naito Foundation to H.O.

## Author contributions

H.O. designed the study; H.O., K.I., and M.K. implemented the study; H.O. prepared crystals and atomic models; and H.O. wrote the report. All authors discussed the results and approved the final version of the manuscript.

## Notes

### Competing Interest Statement

The authors have declared no competing interest.

## References

1. de Bold AJ, Borenstein HB, Veress AT, Sonnenberg H. A rapid and potent natriuretic response to intravenous injection of atrial myocardial extract in rats. Life Sci 28, 89–94 (1981).

2. Currie MG, et al. Bioactive cardiac substances: potent vasorelaxant activity in mammalian atria. Science 221, 71–73 (1983).

3. Grammer RT, Fukumi H, Inagami T, Misono KS. Rat atrial natriuretic factor. Purification and vasorelaxant activity. Biochem Biophys Res Commun 116, 696–703 (1983).

4. Atarashi K, Francosaenz R, Mulrow PJ. Inhibition of Aldosterone Production by Atrial Natriuretic Peptides. Clin Res 32, A785–A785 (1984).

5. Barbee RW, Perry BD, Re RN, Murgo JP, Field LJ. Hemodynamics in Transgenic Mice with Overexpression of Atrial-Natriuretic-Factor. Circulation Research 74, 747–751 (1994).

6. John SW, et al. Genetic decreases in atrial natriuretic peptide and salt-sensitive hypertension. Science 267, 679–681 (1995).

7. Lopez MJ, et al. Salt-Resistant Hypertension in Mice Lacking the Guanylyl Cyclase-a Receptor for Atrial-Natriuretic-Peptide. Nature 378, 65–68 (1995).

8. Oliver PM, et al. Hypertension, cardiac hypertrophy, and sudden death in mice lacking natriuretic peptide receptor A. Proc Natl Acad Sci U S A 94, 14730–14735 (1997).

9. Nojiri T, et al. Atrial natriuretic peptide prevents cancer metastasis through vascular endothelial cells. Proc Natl Acad Sci U S A 112, 4086–4091 (2015).

10. Kangawa K, Matsuo H. Purification and complete amino acid sequence of alpha-human atrial natriuretic polypeptide (alpha-hANP). Biochem Biophys Res Commun 118, 131–139 (1984).

11. Sudoh T, Kangawa K, Minamino N, Matsuo H. A new natriuretic peptide in porcine brain. Nature 332, 78–81 (1988).

12. Sudoh T, Minamino N, Kangawa K, Matsuo H. C-type natriuretic peptide (CNP): a new member of natriuretic peptide family identified in porcine brain. Biochem Biophys Res Commun 168, 863–870 (1990).

13. Nakao K, Ogawa Y, Suga S, Imura H. Molecular biology and biochemistry of the natriuretic peptide system. I: Natriuretic peptides. J Hypertens 10, 907–912 (1992).

14. Koller KJ, et al. Selective activation of the B natriuretic peptide receptor by C-type natriuretic peptide (CNP). Science 252, 120–123 (1991).

15. Nakao K, Ogawa Y, Suga S, Imura H. Molecular biology and biochemistry of the natriuretic peptide system. II: Natriuretic peptide receptors. J Hypertens 10, 1111–1114 (1992).

16. Chusho H, et al. Dwarfism and early death in mice lacking C-type natriuretic peptide. Proc Natl Acad Sci U S A 98, 4016–4021 (2001).

17. Schweitz H, Vigne P, Moinier D, Frelin C, Lazdunski M. A new member of the natriuretic peptide family is present in the venom of the green mamba (Dendroaspis angusticeps). J Biol Chem 267, 13928–13932 (1992).

18. Lisy O, et al. Renal actions of synthetic dendroaspis natriuretic peptide. Kidney Int 56, 502–508 (1999).

19. Best PJ, Burnett JC, Wilson SH, Holmes DR, Jr., Lerman A. Dendroaspis natriuretic peptide relaxes isolated human arteries and veins. Cardiovasc Res 55, 375–384 (2002).

20. Chinkers M, et al. A membrane form of guanylate cyclase is an atrial natriuretic peptide receptor. Nature 338, 78–83 (1989).

21. Garbers DL, et al. Membrane guanylyl cyclase receptors: an update. Trends Endocrinol Metab 17, 251–258 (2006).

22. Potter LR. Guanylyl cyclase structure, function and regulation. Cell Signal 23, 1921–1926 (2011).

23. Ogawa H, Qiu Y, Ogata CM, Misono KS. Crystal structure of hormone-bound atrial natriuretic peptide receptor extracellular domain: rotation mechanism for transmembrane signal transduction. J Biol Chem 279, 28625–28631 (2004).

24. van den Akker F, Zhang X, Miyagi M, Huo X, Misono KS, Yee VC. Structure of the dimerized hormone-binding domain of a guanylyl-cyclase-coupled receptor. Nature 406, 101–104 (2000).

25. Misono KS, Sivasubramanian N, Berkner K, Zhang X. Expression and purification of the extracellular ligand-binding domain of the atrial natriuretic peptide (ANP) receptor: monovalent binding with ANP induces 2:2 complexes. Biochemistry 38, 516–523 (1999).

26. Okamoto N, Ogawa H, Toyoshima C. A new method for establishing stable cell lines and its use for large-scale production of human guanylyl cyclase-B receptor and of the extracellular domain. Biochem Biophys Res Commun 426, 260–265 (2012).

27. Misono KS. Atrial natriuretic factor binding to its receptor is dependent on chloride concentration: A possible feedback-control mechanism in renal salt regulation. Circ Res 86, 1135–1139 (2000).

28. Ogawa H, Qiu Y, Philo JS, Arakawa T, Ogata CM, Misono KS. Reversibly bound chloride in the atrial natriuretic peptide receptor hormone-binding domain: possible allosteric regulation and a conserved structural motif for the chloride-binding site. Protein Sci 19, 544–557 (2010).

29. Singh G, Kuc RE, Maguire JJ, Fidock M, Davenport AP. Novel snake venom ligand dendroaspis natriuretic peptide is selective for natriuretic peptide receptor-A in human heart: downregulation of natriuretic peptide receptor-A in heart failure. Circ Res 99, 183–190 (2006).

30. Willenbrock RC, Tremblay J, Garcia R, Hamet P. Dissociation of Natriuresis and Diuresis and Heterogeneity of the Effector System of Atrial Natriuretic Factor in Rats. J Clin Invest 83, 482–489 (1989).

31. Koyama S, et al. An Oxidized Analog of Alpha-Human Atrial Natriuretic Polypeptide Is a Selective Agonist for the Atrial-Natriuretic-Polypeptide Clearance Receptor Which Lacks a Guanylate-Cyclase. Eur J Biochem 203, 425–432 (1992).

32. Maack T, et al. Physiological role of silent receptors of atrial natriuretic factor. Science 238, 675–678 (1987).

33. Bovy PR. Structure Activity in the Atrial Natriuretic Peptide (Anp) Family. Med Res Rev 10, 115–142 (1990).

34. Schulz S, Green CK, Yuen PS, Garbers DL. Guanylyl cyclase is a heat-stable enterotoxin receptor. Cell 63, 941–948 (1990).

35. Currie MG, et al. Guanylin - an Endogenous Activator of Intestinal Guanylate-Cyclase. P Natl Acad Sci USA 89, 947–951 (1992).

36. Hamra FK, et al. Uroguanylin - Structure and Activity of a 2nd Endogenous Peptide That Stimulates Intestinal Guanylate-Cyclase. P Natl Acad Sci USA 90, 10464–10468 (1993).

37. Field M, Graf LH, Laird WJ, Smith PL. Heat-Stable Enterotoxin of Escherichia-Coli - Invitro Effects on Guanylate Cyclase Activity, Cyclic-Gmp Concentration, and Ion-Transport in Small-Intestine. P Natl Acad Sci USA 75, 2800–2804 (1978).

38. Chen HH, Lainchbury JG, Burnett JC, Jr. Natriuretic peptide receptors and neutral endopeptidase in mediating the renal actions of a new therapeutic synthetic natriuretic peptide dendroaspis natriuretic peptide. J Am Coll Cardiol 40, 1186–1191 (2002).

39. Lisy O, Huntley BK, McCormick DJ, Kurlansky PA, Burnett JC, Jr. Design, synthesis, and actions of a novel chimeric natriuretic peptide: CD-NP. J Am Coll Cardiol 52, 60–68 (2008).

40. Lee CY, et al. Cenderitide: structural requirements for the creation of a novel dual particulate guanylyl cyclase receptor agonist with renal-enhancing in vivo and ex vivo actions. Eur Heart J Cardiovasc Pharmacother 2, 98–105 (2016).

41. Johns DG, et al. Dendroaspis natriuretic peptide binds to the natriuretic peptide clearance receptor. Biochem Bioph Res Co 358, 145–149 (2007).

42. Fuller F, et al. Atrial Natriuretic Peptide Clearance Receptor - Complete Sequence and Functional Expression of Cdna Clones. Journal of Biological Chemistry 263, 9395–9401 (1988).

43. He X, Chow D, Martick MM, Garcia KC. Allosteric activation of a spring-loaded natriuretic peptide receptor dimer by hormone. Science 293, 1657–1662 (2001).

44. He XL, Dukkipati A, Garcia KC. Structural determinants of natriuretic peptide receptor specificity and degeneracy. J Mol Biol 361, 698–714 (2006).

45. Suga SI, et al. Receptor Selectivity of Natriuretic Peptide Family, Atrial-Natriuretic-Peptide, Brain Natriuretic Peptide, and C-Type Natriuretic Peptide. Endocrinology 130, 229–239 (1992).

46. Cohen D, Koh GY, Nikonova LN, Porter JG, Maack T. Molecular determinants of the clearance function of type C receptors of natriuretic peptides. Journal of Biological Chemistry 271, 9863–9869 (1996).

47. Potter LR. Natriuretic peptide metabolism, clearance and degradation. FEBS J 278, 1808–1817 (2011).

48. Nakao K, et al. The pharmacokinetics of alpha-human atrial natriuretic polypeptide in healthy subjects. Eur J Clin Pharmacol 31, 101–103 (1986).

49. Yandle TG, Richards AM, Nicholls MG, Cuneo R, Espiner EA, Livesey JH. Metabolic clearance rate and plasma half life of alpha-human atrial natriuretic peptide in man. Life Sci 38, 1827–1833 (1986).

50. Horst BG, et al. Allosteric activation of the nitric oxide receptor soluble guanylate cyclase mapped by cryo-electron microscopy. Elife 8, (2019).

51. Otwinowski Z, Minor W. Processing of X-ray diffraction data collected in oscillation mode. Method Enzymol 276, 307–326 (1997).

52. Vagin A, Teplyakov A. MOLREP: an automated program for molecular replacement. J Appl Crystallogr 30, 1022–1025 (1997).

53. Brunger AT, et al. Crystallography & NMR system: A new software suite for macromolecular structure determination. Acta Crystallogr D 54, 905–921 (1998).

54. Liebschner D, et al. Macromolecular structure determination using X-rays, neutrons and electrons: recent developments in Phenix. Acta Crystallographica Section D-Structural Biology 75, 861–877 (2019).

