## Supplementary Methods, Supplemental Table 1 for "Structural insight into hormone recognition and transmembrane signaling by the atrial natriuretic peptide receptor"

### Precise modeling of bound ligands

Modeling the bound ligands was highly challenging. Although the bound peptides had no internal symmetry, they showed an obvious two-fold symmetry. These findings indicate that the two ANP molecules have two alternative conformations (orientations) with two-fold symmetry at their binding sites with an occupancy of 50% (Supplementary Fig. 3-c). To resolve difficulties with modeling the bound ligands, $|F_{\mathrm{obs(peptide1)}}|$-$|F_{\mathrm{obs(peptide2)}}|$ electron density maps between multiple ligand complexes were created to detect differences in the specific amino acid sequences (Supplementary Fig. 3d-i). For example, the amino acids differ between human and rat ANP only at position 12, being Met in humans but Ile in rats (Supplementary Fig 3h). A sulfur atom is located at the tip of the Met side chain, so it has a larger electron density than the carbon atom in Ile. Therefore, the electron density should appear at the sulfur site of Met12 in the $|F_{\mathrm{obs(hANP[1\text{-}28])}}| - |F_{\mathrm{obs(rANP[1\text{-}28])}}|$ map. Indeed, the $|F_{\mathrm{obs(hANP)}}|$-$|F_{\mathrm{obs(rANP)}}|$ map at 12σ clearly shows the sulfur site of Met12 in the hANP (or Ile12 in the rANP[1-28]) in the two alternative conformations (Supplementary Fig. 2d). This calculation is possible because the unit cell dimensions of the crystals other than those of rANP[4-17, 23] are essentially the same (Supplementary Table 1). This strategy was also useful for the truncated ligands. Indeed, the $|F_{\mathrm{obs(rANP[1\text{-}28])}}|$-$|F_{\mathrm{obs(rANP[7\text{-}27])}}|$ map at 5σ shows the locations of Tyr28 in the two conformations (Supplementary Fig. 2g, h). By defining the locations of specific side chains using this strategy (Supplementary Figs. 3 – 5), we identified 11 residues, each in ANP and DNP, and modeled amino acid residues 4 – 28 of hANP[1-28]/ or rANP[1-28], and amino acid residues 1 – 28 of DNP with no ambiguity (Fig. 2 a, d, Supplementary Fig. 8).

**Supplemental Table 1. Data collection and refinement statistics**

| **Data collection** | rANP[1-28] | hANP[1-28] | DNP |
|---|---|---|---|
| Space group | $P6_1$ | $P6_1$ | $P6_1$ |
| No. crystals | 2 | 2 | 2 |
| Cell dimensions | | | |
| *a*, *b*, *c* (Å) | 100.2, 100.2, 261.9 | 100.2, 100.2, 261.7 | 99.6, 99.6, 262.5 |
| Resolution (Å) | 50.0-2.45 (2.52-2.45)* | 50.0-2.45 (2.52-2.45)* | 50.0-2.45 (2.52-2.45)* |
| $R_{merge}$ | 6.0 (35.5) | 4.0 (36.7) | 5.3 (25.5) |
| $I/\sigma$ | 30.5 (2.6) | 34.9 (3.2) | 30.0 (4.6) |
| $CC_{1/2}$ | 0.99 (0.56) | 0.99 (0.51) | 0.99 (0.66) |
| Completeness (%) | 99.6 (98.8) | 96.8 (98.1) | 96.3 (99.5) |
| Redundancy | 22.3 (19.5) | 27.8 (19.8) | 22.1 (16.5) |
| **Refinement** | | | |
| Resolution (Å) | 32.9-2.45 (2.53-2.45) | 43.5-2.45 (2.53-2.45) | 30.7-2.45 (2.53-2.45) |
| No. reflections | 52,690 (4,850) | 52,727 (4,831) | 51,983 (4,880) |
| $R_{work}$/ $R_{free}$ | 0.193 / 0.249 (0.306 / 0.314) | 0.205 / 0.259 (0.305 / 0.335) | 0.201 / 0.241 (0.290 / 0.312) |
| No. atoms | | | |
| Protein | 6,944 | 6,944 | 6,944 |
| Ligand | 189 | 189 | 231 |
| Waters, ions | 135 | 133 | 127 |
| *B*-factors | | | |
| Protein | 89.9 | 86.1 | 63.3 |
| Ligand | 86.4 | 80.9 | 58.0 |
| Waters, ions | 68.3 | 64.4 | 42.3 |
| R.m.s deviations | | | |
| Bond lengths (Å) | 0.006 | 0.009 | 0.006 |
| Bond angles (°) | 0.785 | 0.994 | 0.829 |

*Highest resolution shell is shown in parenthesis.

| **Data collection** | hANP[7-28] | hANP[5-27] | rANP[4-17,23] |
|---|---|---|---|
| Space group | $P6_1$ | $P6_1$ | $P6_1$ |
| No. crystals | 2 | 2 | 2 |
| Cell dimensions | | | |
| *a*, *b*, *c* (Å) | 99.8, 99.8, 260.4 | 99.4, 99.4, 259.7 | 100.2, 100.2, 270.4 |
| Resolution (Å) | 50.0-2.7 (2.78-2.70)* | 50.0-2.85 (2.93-2.85)* | 50.0-3.2 (3.29-3.20)* |
| $R_{merge}$ | 3.1 (30.7) | 7.8 (24.3) | 4.8 (37.3) |
| $I/\sigma$ | 23.5 (3.5) | 26.4 (5.3) | 27.6 (3.5) |
| $CC_{1/2}$ | 0.99 (0.53) | 0.99 (0.55) | 0.99 (0.42) |
| Completeness (%) | 99.3 (99.5) | 90.2 (99.9) | 97.4 (99.4) |
| Redundancy | 10.9 (9.6) | 15.4 (7.3) | 23.1 (19.7) |
| **Refinement** | | | |
| Resolution (Å) | 43.3-2.70 (2.81-2.70) | 43.1-2.85 (3.03-2.85) | 33.5-3.20 (3.45-3.20) |
| No. reflections | 39,898 (4,415) | 30,550 (5,626) | 24,970 (5,024) |
| $R_{work}/ R_{free}$ | 0.190 / 0.234 (0.314 / 0.370) | 0.173 / 0.230 (0.267 / 0.312) | 0.198 (0.237) (0.307/0.396) |
| No. atoms | | | |
| Protein | 6,944 | 6,944 | 6,944 |
| Ligand | 166 | 166 | 110 |
| Waters, ions | 131 | 103 | 2 |
| B-factors | | | |
| Protein | 89.1 | 91.8 | 116.8 |
| Ligand | 101.1 | 91.2 | 124.1 |
| Waters, ions | 65.6 | 68.8 | 78.7 |
| R.m.s deviations | | | |
| Bond lengths (Å) | 0.009 | 0.008 | 0.007 |
| Bond angles (º) | 0.989 | 0.965 | 0.822 |

```
rANP[1-28]    :     SLRRSSCFGGRIDRIGAQSGLGCNSFRY
hANP[1-28]    :     SLRRSSCFGGRMDRIGAQSGLGCNSFRY
DNP           :     EVKYDPCFGHKIDRINHVSNLGCPSLRDPRPNAPSTSA

hBNP          :SPKMVQGSGCFGRKMDRISSSSGLGCKVLRRH
hCNP          :      GLSKGCFGLKIDRIGSMSGLGCN
```

**Supplementary Fig. 1. Sequence alignment of the natriuretic peptides.** Upper panel, natriuretic peptides determined in this study. Two Cys residues (blue with underscore) form SS-bond, and peptide forms a cyclic structure. Residues that contribute to forming pseudo two-fold symmetry in tertiary structure are boxed and linked with lines. Lower panel, other natriuretic peptide families with undetermined structures. Residues that might contribute to forming pseudo two-fold symmetry in tertiary structure are boxed and linked with lines.

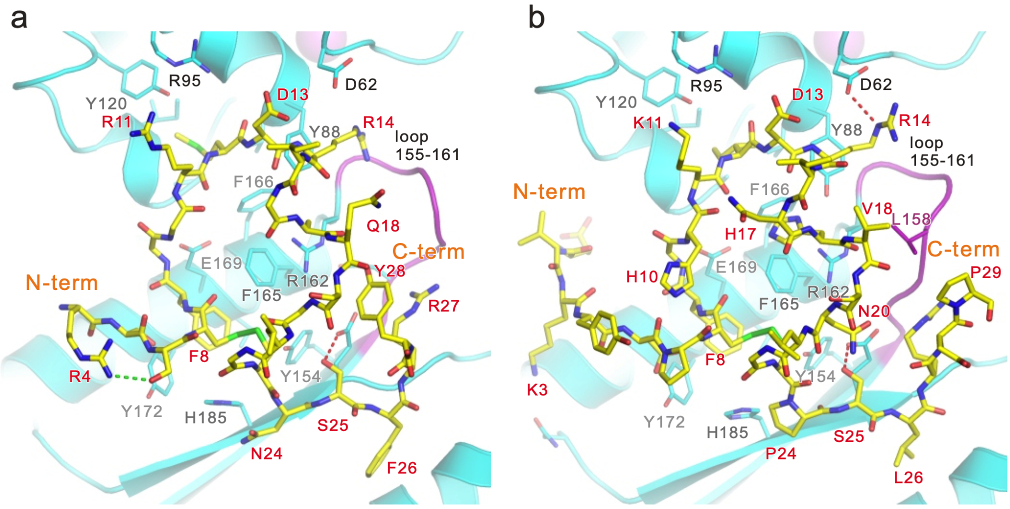

**Supplementary Fig. 2. Structures of the bound peptides. Magnified views of bound hANP and DNP looking from monomer A.** (a) hANP. (b) DNP. Induced fit occurred at loop 155 – 161.

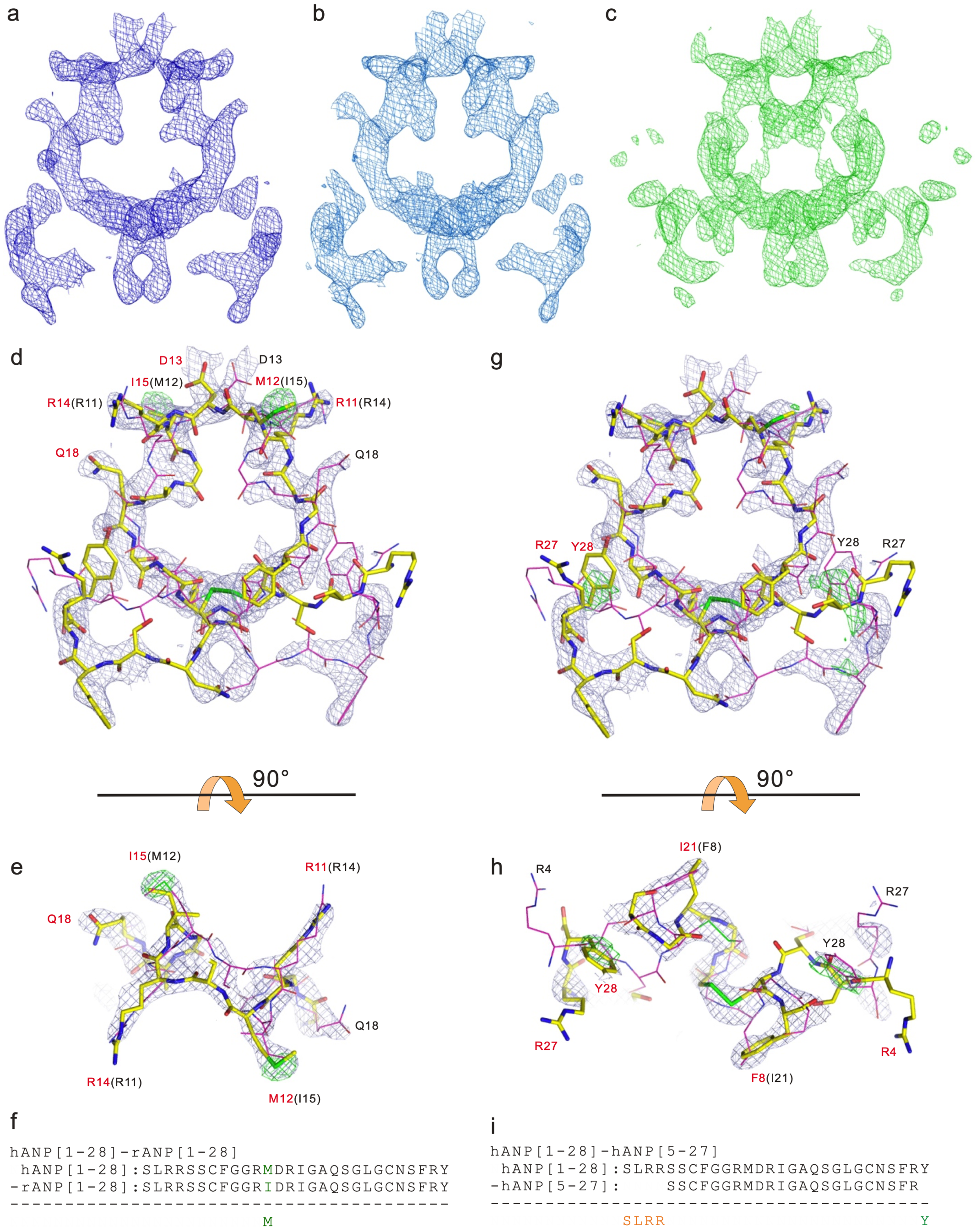

**Supplementary Fig. 3. Electron density maps around bound peptides.** Initial electron density maps around (a) rANP[1-28], (b) hANP[1-28], (c) DNP contour level at 2.5σ. (d–g) $|F_{obs(peptide1)}|$-$|F_{obs(peptide2)}|$ maps between multiple ligand complexes. (d, e, f) $|F_{obs(hANP[1-28])}|$-$|F_{obs(rANP[1-28])}|$ map at 12.5σ. The map clearly shows the sulfur site of Met12 in hANP[1-28]. (g, h, i) $|F_{obs(rANP[1-28])}|$-$|F_{obs(rANP[7-27])}|$ map at 5.0σ. The map clearly shows locations of Tyr28. Since two ANP molecules are comprised of two ligand molecules having two alternative conformations (orientations) with two-fold symmetry at their binding sites with an occupancy of 50%, one conformation is shown as a yellow stick and the other is shown as a thin purple stick.

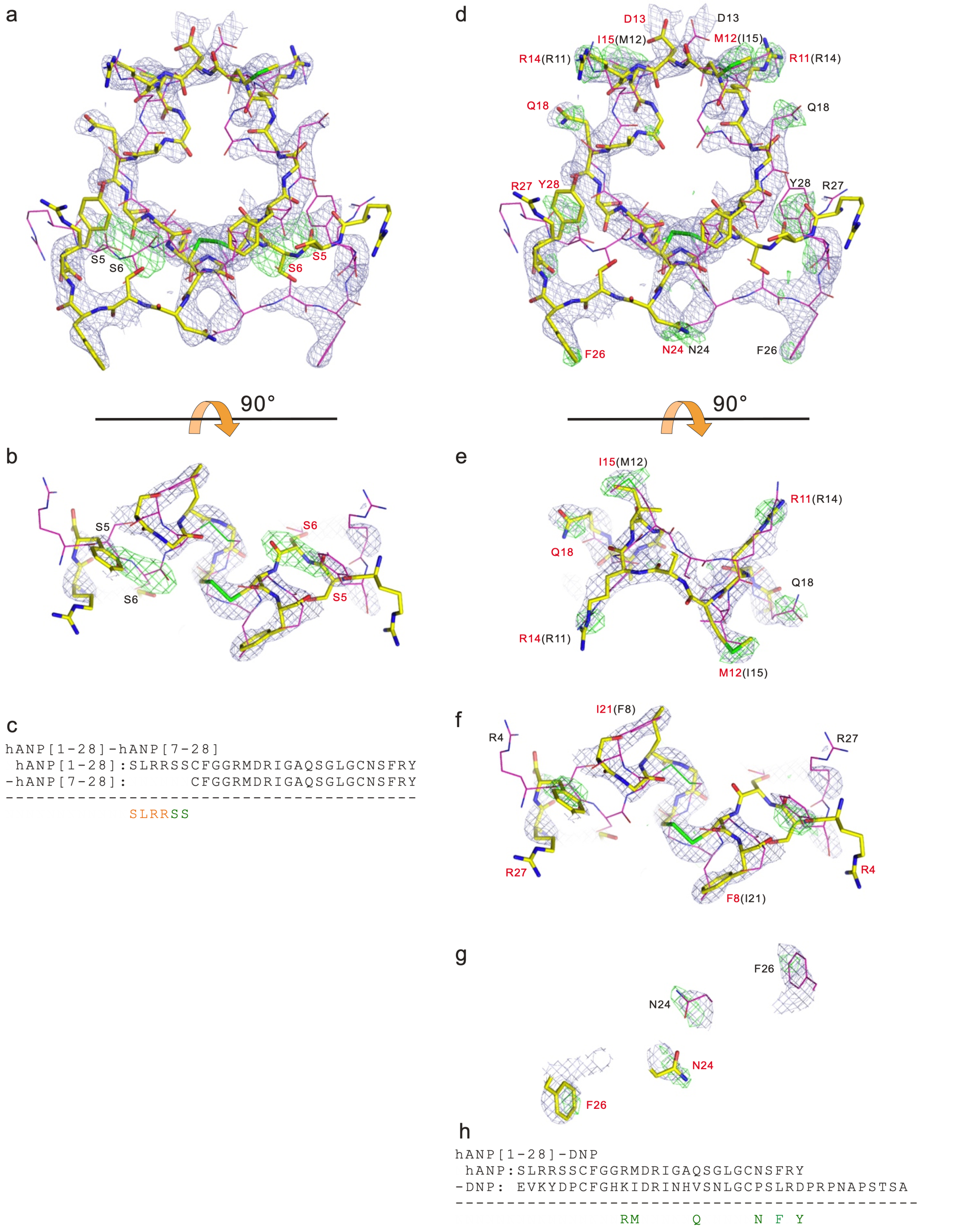

**Supplementary Fig. 4. $|F_{obs}|$-$|F_{obs}|$ electron density maps around the bound peptides.** (a, b, c) $|F_{obs(hANP[1\text{-}28])}|$-$|F_{obs(hANP[7\text{-}28])}|$ map at 5.0σ. (d, e, f, g, h) $|F_{obs(hANP[1\text{-}28])}|$-$|F_{obs(DNP)}|$ map at 4.0σ. The map clearly shows the locations of Arg11, Met12, Gln18, Asn24, Phe26, and Tyr28 in hANP[1-28]. Since two ANP molecules are comprised of two alternative conformations (orientations) with two-fold symmetry at their binding sites with an occupancy of 50%, one conformation is shown as a yellow stick and the other is shown as a thin purple stick.

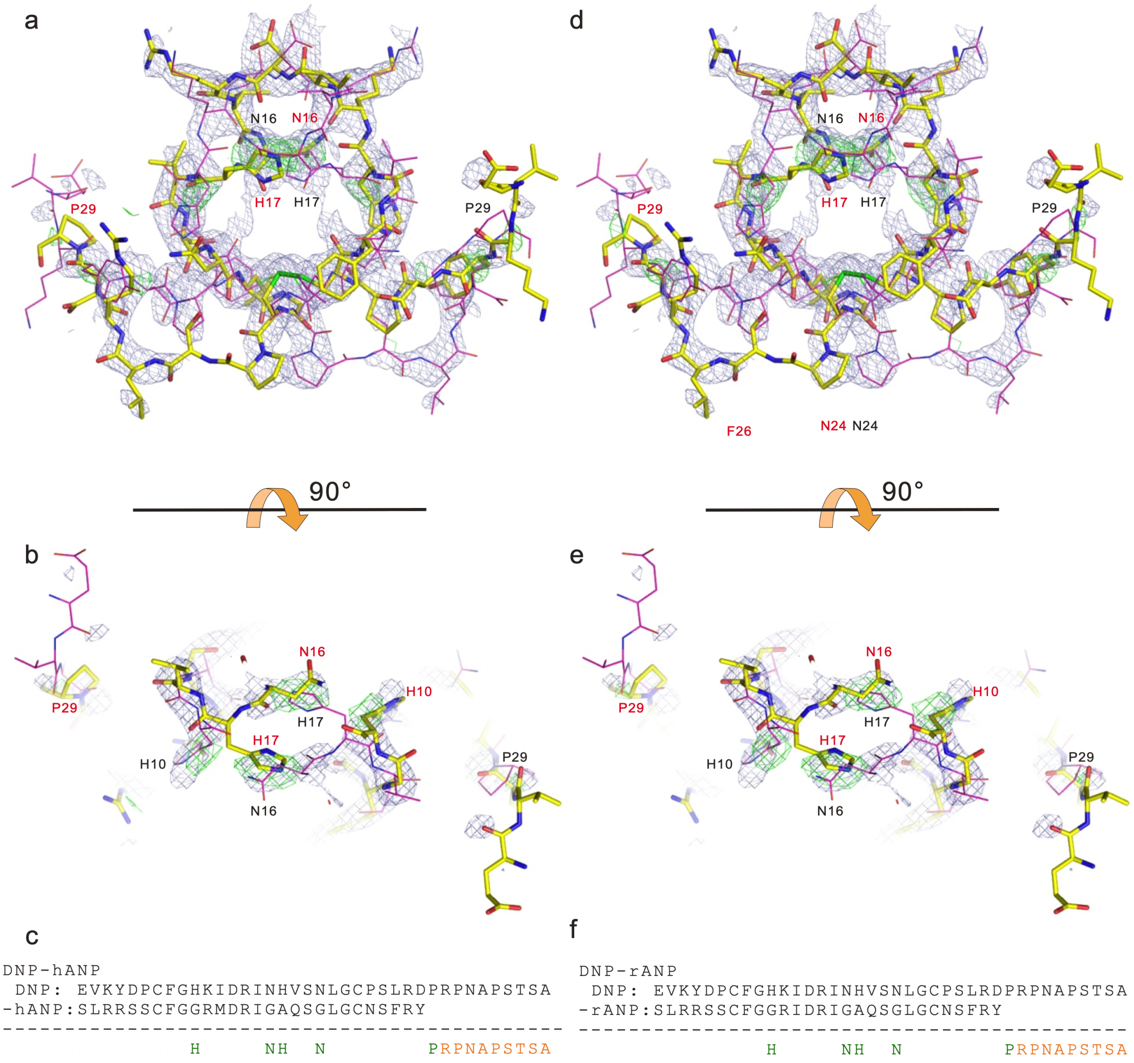

**Supplementary Fig. 5. $|F_{obs}|$-$|F_{obs}|$ electron density maps around the bound peptides.** (a, b, c) $|F_{obs(DNP)}|$-$|F_{obs(hANP(1\text{-}28))}|$ map at 4.5σ. The map clearly shows the locations of His10, Asn16, His17, Asn20 and Pro29 in the DNP. (d, e, f) $f_{o(DNP)}$-$f_{o(rANP(1\text{-}28))}$ map at 4.5σ. The map clearly shows the locations of His10, Asn16, His17, Asn20 and Pro29 in the DNP. Since two DNP molecules are comprised of two alternative conformations (orientations) with two-fold symmetry at their binding sites with an occupancy of 50%, one conformation is shown as a yellow stick and the other is shown as thin a purple stick.

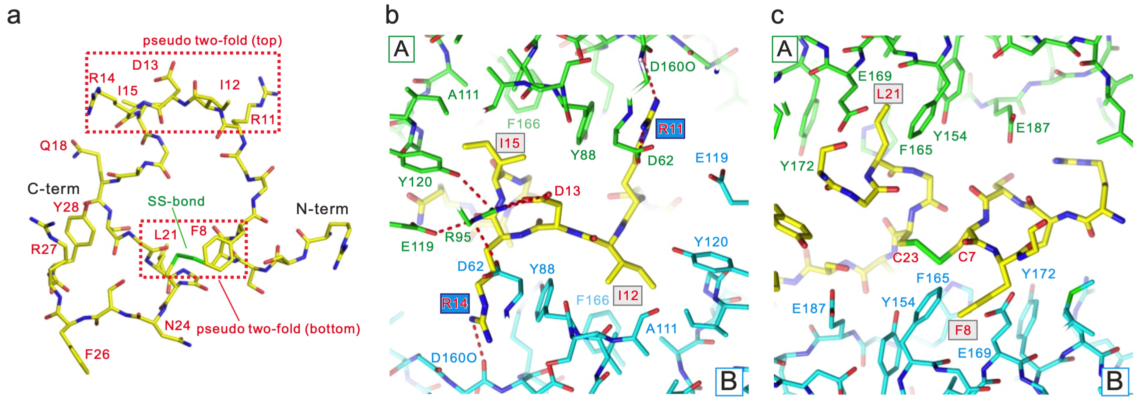

**Supplementary Fig. 6. Pseudo two-fold regions of bound peptides.** (a) The structure of rANP[1-28] bound to the ANPR. rANP[1-28] have pseudo-two-fold symmetry at the top and bottom of ring structures, and both regions are surrounded by red dotted boxes. Magnified views of the pseudo two-fold symmetry region at the top (b) and bottom (c) of the ring in rANP[1-28]. All molecules are represented as sticks. Monomers A and B are green and cyan, respectively. The bound ligands are shown as yellow sticks. Red, oxygen; blue, nitrogen; yellow-green, sulfite atoms. Red dashed lines represent hydrogen bonds. (b) Two independent pairs with pseudo-two-fold symmetry at the top of each ring structure are centered on Asp13. One is hydrophobic and consists of Ile12 and Ile15, whereas the other is hydrophilic and consists of Arg11 and Arg14. Both pairs can enter symmetrical hydrophobic or hydrophilic pockets of ANPR monomers. (c) Hydrophobic pairs of Phe8 and Leu21 centered on the disulfide bond composed of Cys7 and Cys23 have pseudo two-fold symmetry. Hydrophobic pairs can enter symmetrical hydrophobic pocket of ANPR monomers.

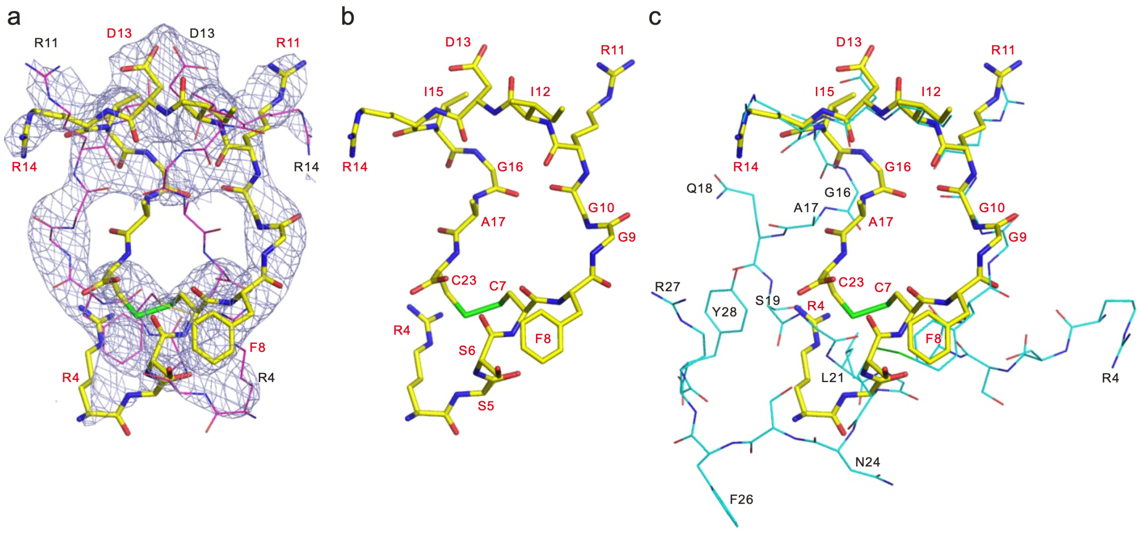

**Supplementary Fig. 7. Structure of bound rANP[4-17, 23].** (a) Initial electron density map around rANP[4-17, 23] at 2.5σ. (b) rANP[4-17, 23] molecule. (c) Comparison with rANP[1-28] shown as thin cyan stick. Since two rANP[4-17, 23] molecules have two alternative conformations (orientations) with two-fold symmetry at their binding sites with an occupancy of 50%, one conformation is shown as a yellow stick and another is shown as a thin purple stick.

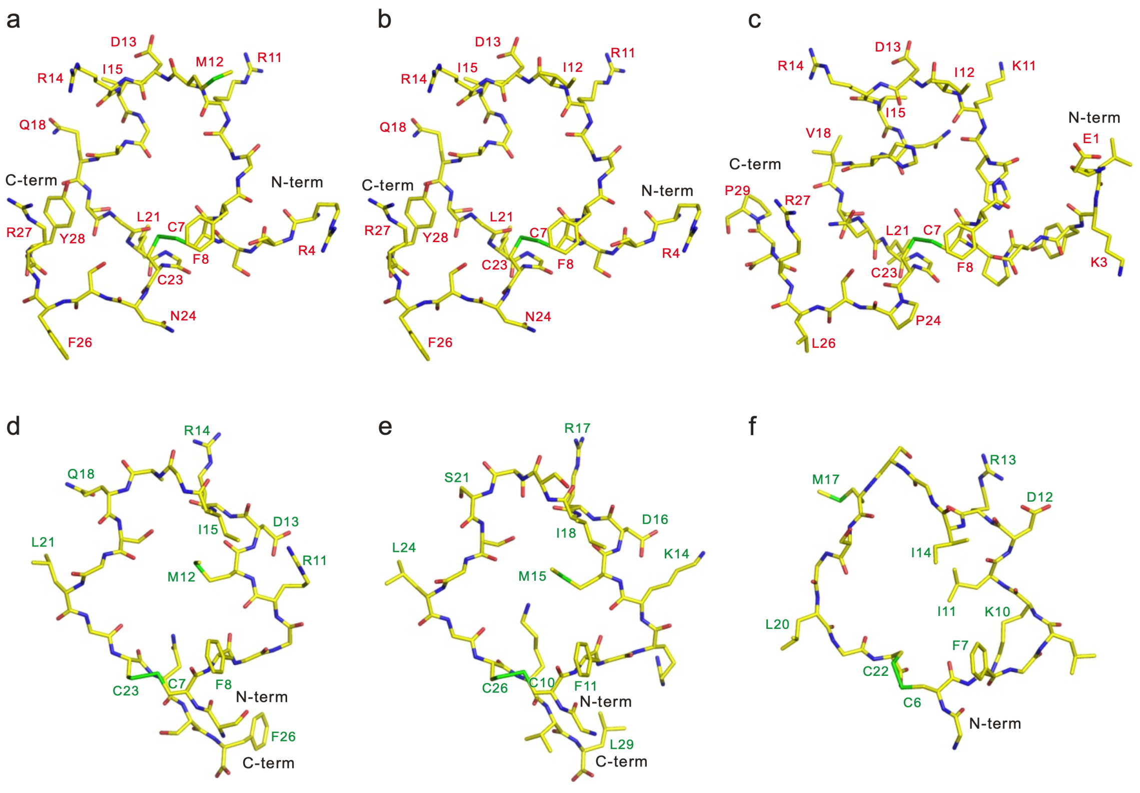

**Supplementary Fig. 8. Comparison with natriuretic peptides bound to the NPRC.** (a-c) Natriuretic peptides complexed with the ANPR. (a) hANP[1-28], (b) rANP[1-28] and (c) DNP. The ring structures of the peptides are similar. They possess pseudo two-fold symmetry at the top and bottom region of their ring structures. (d-f) Natriuretic peptides complexed with the NPRC. (d) hANP[1-28], (b) hBNP and (c) hCNP. The structures of the bound peptides do not possess pseudo two-fold symmetry in the ring structures, and they are extremely different from the peptides complexed with the ANPR.
